# The complete mitochondrial genome of *Heikeopsis japonica* (Decapoda: Brachyura: Dorippidae): Phylogenetic implications of the family Dorippidae

**DOI:** 10.1101/2024.08.21.609022

**Authors:** Xiaoke Pang, Yifan Zhao, Yufei Liu, Xueqiang Lu

**Author notes:** Corresponding author. Xueqiang Lu.

## Abstract

Characterizing complete mitochondrial genome (mitogenome) supports comprehensive understanding in gene arrangement, molecular evolution and phylogenetic status. Previously, no studies have been conducted on the complete mitogenomes of the Dorippidae family within infraorder Brachyura. Herein, we firstly determined the sequence of *Heikeopsis japonica* (Decapoda: Brachyura: Dorippidae) mitogenome by high-throughput sequencing. Its mitogenome sequence is 15,980 bp in size, with 13 protein-coding genes, two ribosomal RNA genes, and 22 transfer RNA genes. The genome has a high A + T content of 73.52%, and low G + C content of 26.48%. The AT-skew is negative, -0.0079; and the GC-skew is positive, 0.322. The phylogenetic analysis of *H. japonica* using 40 Brachyuran mitogenome datasets indicated that *Pyrhila pisum* of family Leucosiidae had the closet relationship with *H. japonica*. Further selection pressure analysis of *H. japonica* suggested the presence of strong positive purification selection in Brachyuran. Meanwhile, a total of 31 residues located in genes *atp6, atp8, cox1*-*3, cob, nad1*-*5*, and *nad4L* were detected as the positively selected sites. This study reports the first available complete mitogenome of family Dorippidae, and our results could offer a useful phylogenetic implication of Dorippidae in the infraorder Brachyura.

## 1. Introduction

Mitochondria, which carry important genetic information and possess a relatively independent genetic system, have always been a hotspot in the research area of molecular biology (Martijn *et al*. 2018). Mitochondrial DNA is maternally inherited, with easily accessible feature, small genome size, conservative gene organization and considerable mutation rate (Clayton 2000; Bernt *et al*. 2013). Mitochondrial genomes (mitogenomes) usually offer more detailed genetic information than partial mitochondrial genes and possess the advantages of distinctive evolutionary rates between several segments (Sun *et al*. 2018a; Li *et al*. 2019). With the rapid development of high-throughput sequencing technology, mitogenomes are increasingly used in genetic evolution, phylogenetic analysis and molecular adaptive mechanisms (Kou *et al*. 2020; Zhong *et al*. 2020). Generally speaking, the lengths of metazoan genome sequences are 15-20 kb. It usually comprises 13 protein coding genes (PCGs), two ribosomal RNA (rRNA) genes, and 22 transfer RNA (tRNA) genes. The structural features and gene arrangements of the genetic sequence presented by mitogenome usually vary in each species (Kundu *et al*. 2019). Therefore, mitogenomic composition can be characterized for investigating the phylogenetic analysis and adaptative evolution of new species (Feng *et al*. 2021; Wang *et al*. 2023).

Dorippoid crabs (superfamily Dorippoidea MacLeay, 1838) are an important part of macrobenthic invertebrates in the subtidal zone of marine ecosystems. These species are critical in the degradation of organic substances and the flow of energy in the marine food web (Constable 1999; Carvalho *et al*. 2007). Dorippoid crabs usually use a pair of pereiopods to carry a variety of subjects on the seabed, for example, shell crater fragments, sea urchins, wood chips and leaves for protective concealment (Ng and Rahayu 2002; Castro 2005). Thus, a lot of organic substances are successfully transferred among different species. Dorippoids are divided into three families: Dorippidae MacLeay 1838, Ethusidae Guinot 1977, and Orithyiidae Dana 1852. However, only one mitogenome of these species (*Orithyia sinica*, family Orithyiidae) has been reported currently. Dorippid crabs (family Dorippidae MacLeay, 1838) have important ecological value in marine ecosystem, which are proved to be an essential feeding source or companion organism of many important fishery species except for their transferring function (Zairion *et al*. 2017). Most of the previous studies on family Dorippidae were conducted based on the distribution of the species, such as their records in Indonesia (Ng and Rahayu 2002), Iraq (Yasser and Naser 2019), Philippines (Takeda and Manuel-Santos 2006) and India (Devi and Kumar 2017). Studies on the mitochondrial genes of dorippid crabs in the family Dorippidae, are relatively rare and none of any complete species mitogenome of Dorippidae has been reported up to date.

In this study, we sequenced the first complete mitogenome of *H. japonica* that belongs to the family Dorippidae. The genome structure, nucleotide composition, and gene order of *H. japonica* were characterized. Furthermore, we reconstructed a phylogenetic tree to reveal the evolutionary status of *H. japonica* within infraorder Brachyura as well as analyzed the natural selection pressure. Integrally, our study provides the first mitogenome data of family Dorippidae, and offers an elementary understanding of its phylogenetic position and evolutionary journey. This is of great significance for the supplement of molecular data and the management of the biological resources of crabs in marine ecosystems.

## 2. Materials and methods

### 2.1 Sample preparation and DNA extraction

An individual specimen of *H. japonica* was collected from Bohai Bay, China in August 2022. Morphological identification at species level was carried out according to the main idea of Sin et al (Sin *et al*. 2009). The specimen was stored in 95% ethanol until DNA extraction. DNeasy tissue kit (Qiagen, Beijing, China) was used to extract the total genomic DNA from the muscle tissues of *H. japonica*. The collected DNA was preserved at -20°C until the sequencing procedures were performed.

### 2.2 Sequencing, assembly and annotation

After DNA isolation, 1.0 μg of purified DNA fragments were used to conduct paired-end libraries with insert sizes of 450 bp. Sequencing process was carried out by Illumina NovaSeq 6000 at Shanghai BIOZERON Co., Ltd. (Shanghai, China). Before assembly, the raw reads were filtered to obtain high quality reads firstly. We used SOAPdenovo2.04 to assembly these reads into contigs (Luo *et al*. 2012). Then, assembled contigs were aligned to the reference mitogenome (NC_046825.1) via BLAST. Aligned contigs with similarity and query coverage over 80% were arranged with the reference mitogenome as a template. Finally, these reads were aligned to assembled circular genome sketches to cross-check for faulty bases and vacancies were packed with GapFiller v2.1.1.

Mitochondrial genes of *H. japonica* were predicted by MITOS. The position of every PCG was detected by BLAST searches. As for start and stop codons in PCGs, their manual corrections were conducted on SnapGene Viewer according to the reference mitogenomes. The graphic genome structure of *H. japonica* was conducted with CGview tool. The nucleotide composition and codon usage were obtained via EMBOSS v6.6.0.0. The skew analysis was conducted by: AT skew = (A-T)/(A+T) and GC skew = (G-C)/(G+C) (Xia 2013). A, T, G and C represent the occurrences of the nucleotides, respectively. The mitogenome sequences of *H. japonica* were available in GenBank with OQ 434093.

### 2.3 Phylogenetic analysis

Phylogeny of the infraorder Brachyura containing 40 Brachyuran species and *H. japonica* (Table 1) was reconstructed. In addition, species *Alpheus distinguendus* and *Alpheus japonicus* were used as outgroups. The data set was constructed using the sequences of 13 PCGs (*atp6, atp8, cox1-3, cob, nad1-6 and nad4L*), and these sequences were subjected to multiple alignment with MUSCLE 3.8.31. Blurry alignment regions and vacancies in alignment sequences were removed with Gblocks server 0.91b. Model LG + I + G + F was used as the evolutionary model for phylogenetic analysis. Maximum Likelihood (ML) analysis is a typical method for genetic phylogeny in the molecular biology (Nguyen *et al*. 2015). Thus, ML method was used to reconstruct the phylogenetic tree of Brachyura by PhyML v3.0 in this study. The topological robustness of ML analysis was assessed by100 replicates of bootstrap support. Finally, the phylogenetic tree was graphically edited using iTOL 3.4.3.

**Table 1.**
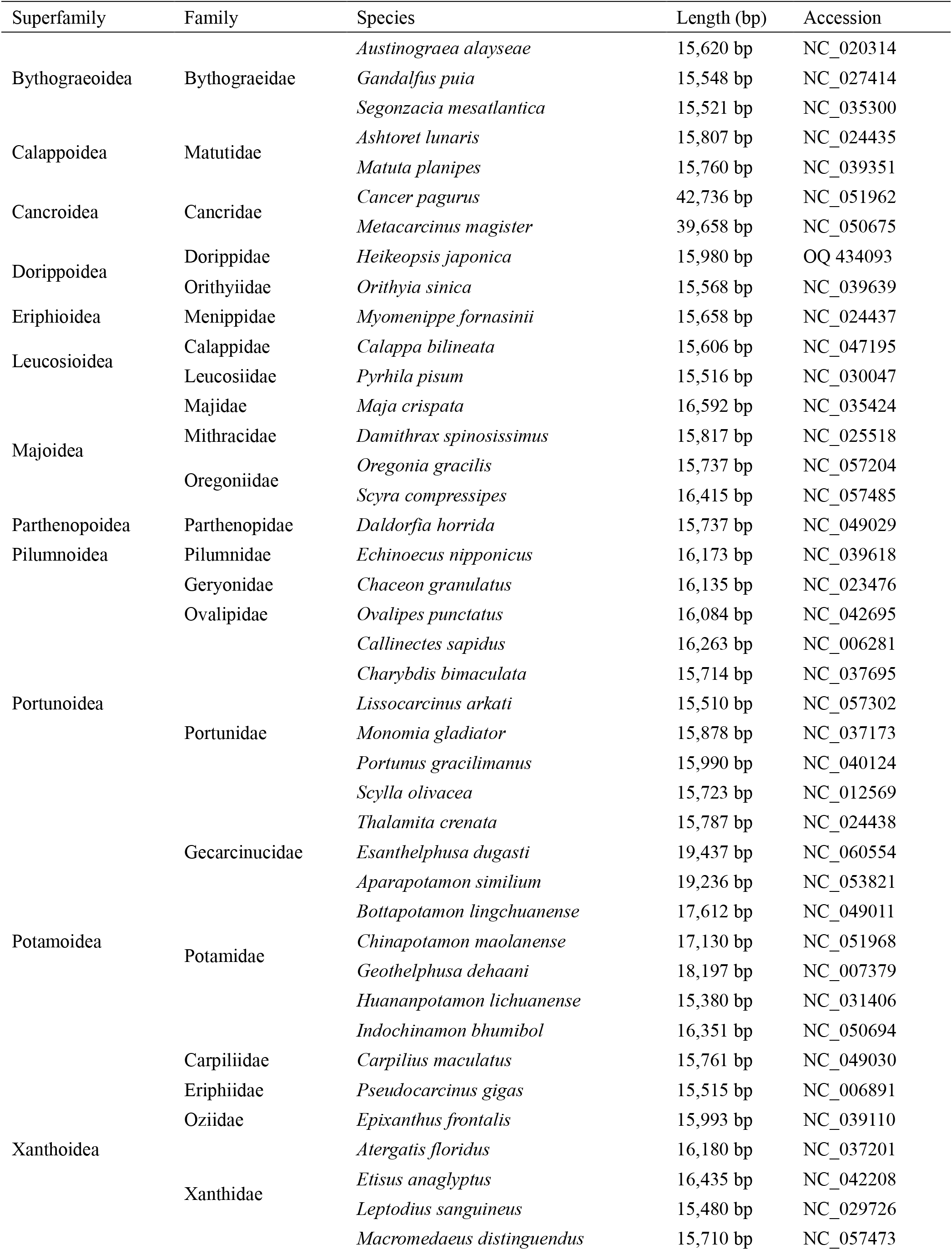
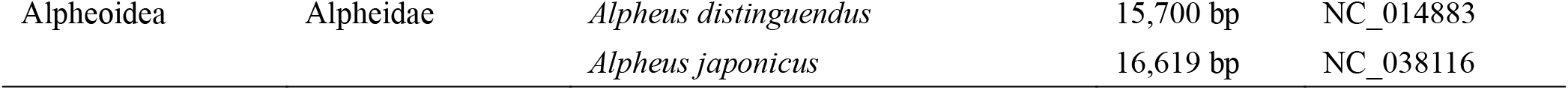
The Brachyuran species used in the phylogenetic tree.

### 2.4 Natural selection analysis

Nonsynonymous and synonymous substitutions (dN and dS) are two important indexes in the species genetic evolution (Yang 1998). The ratio, ω= dN/dS, is a potent value assessing the influence of natural selection force on genetic evolution. When the encoding genes experience positive selection, ω >1. Neutrality and negative selection are denoted by values = 1 and <1, respectively (Yang 1998). Here, all of the 13 PCGs of *H. japonica* were used for selection pressure analyses, and CodeML analysis was used to quantify the ω ratio.

Positive selection analysis of *H. japonica* within infraorder Brachyura were completed with Branch models. Firstly, the ω values were calculated by one-ratio model (M0) in the hypothesis that no adaptive evolution occurs within the genome sequences. This model assumes that all branches of the phylogeny have a single ω ratio (Yang *et al*. 2000). Then, two-ratio model (M2) was implied to determine the selection pressure exerted on target branches. This model allows the background lineages own distinctive ω ratio values compared with the foreground (ω0 and ω1, respectively) (Nielsen and Yang 1998). Thirdly, free-ratio model (M1) was conducted to calculate ω value of each branch by allowing ω ratio between branches to change (Yang *et al*. 2005). M0 and M1 were performed for comparison to confirm whether each Brachyura branch had specific ω ratio. Likelihood ratio tests (LRTs) were used to compare the models with Chi square distribution values. Estimated test statistics were twice the log-likelihood difference (2ΔL). Meanwhile, estimated degrees of freedom in the model used the number differences of parameters.

Finally, the branch site model was fitted to check the positive selection in infraorder Brachyura, which allowed ω value to vary in the protein sites. Potential selected sites within Brachyuran (foreground) lineage were identified using Branch-site model A. The significance levels were detected by model A and null model with ω=1, so that the selection sites in the Brachyuran lineages can be directly determined. It is revealed that model A is fitted more favorable than the corresponding null model when ω value was bigger than 1. Posterior probabilities were computed using Bayes analysis, thus identifying the selection sites on the Brachyuran branch with significant LRTs (Xu *et al*. 2010).

## 3. Results and discussion

### 3.1 *H. japonica* mitogenome sequencing and assembly

The Illumina sequencing data of total genomic DNA was used to sequence and assemble the complete mitogenome of *H. japonica*. This could establish a foundation for the future genetic analysis of the family Dorippidae. A total of 31,257,532 raw data (Q30 = 87.25%) and 29,945,826 bp clean data (Q30 = 88.85%) were obtained. The complete mitogenome of *H. japonica* (OQ 434093) forms a circular molecule, in size of 15,980 bp (Fig. 1; Table 2). It contains 13 PCGs, two rRNA genes and 22 tRNA genes. 23 genes locate in the heavy strand (H), and 14 genes locate in the light strand (L) (Table 2). As reported, this distribution often occurs in the Brachyura crabs’ mitogenomes (Jennings *et al*. 2021; Lu *et al*. 2022). The nucleotide composition of *H. japonica* mitogenome was 36.32% A, 17.5% C, 8.98% G, and 36.9% T, with total AT and GC content of 73.52% and 26.48%, respectively. AT and GC skews are measures of component asymmetry (Chen *et al*. 2021). For *H. japonica*, the AT skew is -0.0079, and the GC skew is 0.3218. The high GC skew value indicates that the nucleotide composition is significantly biased towards G. In addition, there are eight overlaps in the *H. japonica* mitogenome genes, with lengths ranging from 1 to 5 bp (*trnF-ttc*/*nad5, nad4*/*nad4l, trnV-gta*/*rrnS, trnM-atg*/*trnC-tgc, cox1*/*cox2, cox2*/*atp8, atp8*/*atp6, atp6*/*cox3, nad3*/*trnW-tga*).

**Table 2.** Mitogenome organization of *H. japonica*.

| Gene | Strand | Position | Size (bp) | Intergenic spacer | Start codon | Stop codon | Anticodons |
| --- | --- | --- | --- | --- | --- | --- | --- |
| *trnN-aac* | H | 770-834 | 65 | - | - | - | AAC |
| *trnL1-cta* | L | 839-904 | 66 | 4 | - | - | CUA |
| *trnP-cca* | L | 907-970 | 64 | 2 | - | - | CCA |
| *trnY-tac* | L | 985-1050 | 66 | 14 | - | - | UAC |
| *nad1* | L | 1051-1977 | 927 | 0 | ATG | TAA |  |
| *trnH-cac* | L | 1997-2061 | 65 | 19 | - | - | CAC |
| *trnF-ttc* | L | 2062-2125 | 64 | 0 | - | - | UUC |
| *nad5* | L | 2125-3846 | 1722 | -1 | ATT | TAA |  |
| *nad4* | L | >3855-5181 | 1327 | 8 | ATG | A |  |
| *nad4l* | L | 5175-5456 | 282 | -7 | ATA | TAA |  |
| *trnT-aca* | H | 5471-5534 | 64 | 14 | - | - | ACA |
| *cob* | H | 5585-6706 | 1122 | 50 | ATC | TAA |  |
| *trnS2-tca* | H | 6712-6780 | 69 | 5 | - | - | UCA |
| *rrnL* | L | 6816-7860 | 1045 | 35 | - | - |  |
| *trnV-gta* | L | 8055-8119 | 65 | 194 | - | - | GUA |
| *rrnS* | L | 8117-8880 | 764 | -3 | - | - |  |
| *trnQ-caa* | L | 8934-9001 | 68 | 53 | - | - | CAA |
| *trnM-atg* | H | 9018-9082 | 65 | 16 | - | - | AUG |
| *trnC-tgc* | L | 9082-9148 | 67 | -1 | - | - | UGC |
| *cox1* | H | 9150-10,685 | 1536 | 1 | ATG | TAA |  |
| *cox2* | H | 10,685-11,380 | 696 | -1 | ATG | TAA |  |
| *atp8* | H | 11,380-11,541 | 162 | -1 | ATC | TAA |  |
| *atp6* | H | 11,535-12,206 | 672 | -7 | ATA | TAA |  |
| *cox3* | H | 12,206-13,018 | 813 | -1 | ATG | TAA |  |
| *nad3* | H | 13,044-13,388 | 345 | 25 | ATT | TAG |  |
| *trnW-tga* | H | 13,387-13,455 | 69 | -2 | - | - | UGA |
| *trnL2-tta* | H | 13,456-13,520 | 65 | 0 | - | - | UUA |
| *trnI-atc* | H | 13,526-13,590 | 65 | 5 | - | - | AUC |
| *nad2* | H | 13,605->14,562 | 958 | 14 | ATA | T |  |
| *trnK-aaa* | H | 14,563-14,628 | 66 | 0 | - | - | AAA |
| *trnD-gac* | H | 14,629-14,690 | 62 | 0 | - | - | GAC |
| *trnR-cga* | H | 14,691-14,758 | 68 | 0 | - | - | CGA |
| *trnS1-aga* | H | 14,759-14,818 | 60 | 0 | - | - | AGA |
| *nad6* | H | 14,819-15,313 | 495 | 0 | ATG | TAA |  |
| *trnG-gga* | H | 15,380-15,444 | 65 | 66 | - | - | GGA |
| *trnA-gca* | H | 15,446-15,508 | 63 | 1 | - | - | GCA |
| *trnE-gaa* | H | 15,518-15,583 | 66 | 9 | - | - | GAA |

**Fig. 1.**
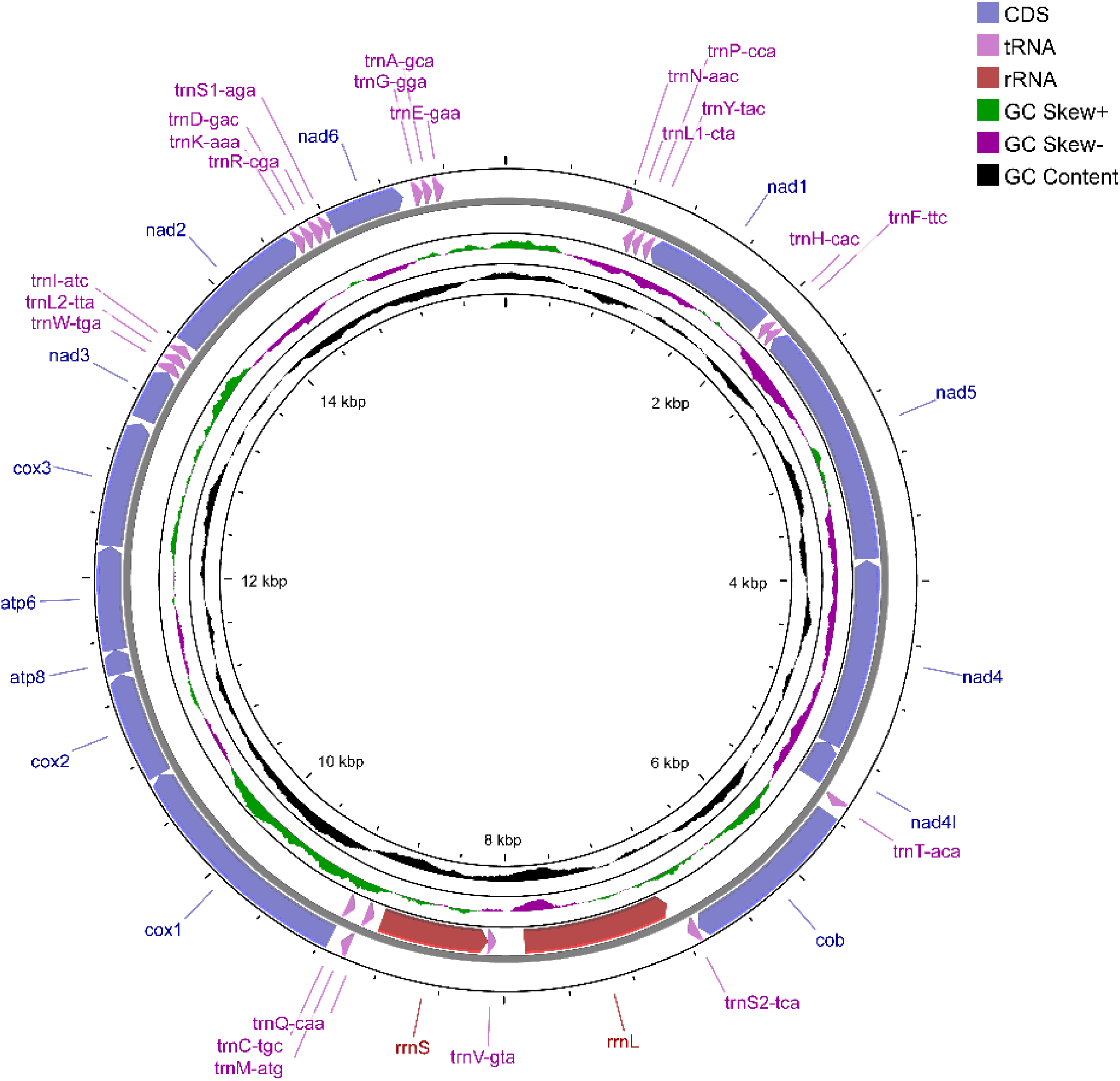
Complete mitogenome map of *H. japonica*. 23 genes are encoded on the heavy (H) strand and 14 genes are encoded on the light (L) strand. Genes for proteins and rRNAs are represented with standard abbreviations. Genes for tRNAs are shown in a single letter for the corresponding amino acid, with two leucine tRNAs and two serine tRNAs distinguished by numerals.

Unique arrangement of mitochondrial genes offers effective phylogenetic signals for investigating the mitochondrial evolution of species (Tan *et al*. 2018). Several studies have confirmed that the gene order in metazoan mitogenomes are conserved, and the presence of gene rearrangements tend to be rare (Basso *et al*. 2017; Wang *et al*. 2018). The gene order of *H. japonica* is different from most of the previously reported co-ordered species with mitogenomes in NCBI, for example, *Calappa bilineata, Chaceon granulates* and *Diogenes edwardsii* (Lu *et al*. 2020; Zhang *et al*. 2020; Pang *et al*. 2023). Through comparative analysis with the decapod ancestor (Lu *et al*. 2022), it was found that *H. japonica* presented a *trnH-cac* translocation, and *trnH-cac* was transferred to *nad1* and *trnF-ttc*, rather than the position between *nad5* and *nad4* as always. Such phenomenon has also occurred in several other crustaceans (Kennish and Williams 1997; Guan *et al*. 2018; Wang *et al*. 2020).

### 3.2 Protein-coding genes

The length of PCGs of *H. japonica* is 11, 057 bp totally, occupying 69.19% of the mitogenome (Table 3). Nine PCGs (*cob, cox1*-*3, atp6, atp8, nad2-3*, and *nad6*) located in the heavy strand, and the rest four genes (*nad1, nad4-5*, and *nad4l*) located in the light strand. Three genes (*nad4L, atp6*, and *nad2*) used ATA as start codon, and two genes (*nad5* and *nad3*) used ATT as start codon, and two genes (*cob* and *atp8*) used ATC. The ATG was determined in the remaining six genes. As for stop codons, the *nad3* gene used TAG, and the most commonly used codon TAA was found in ten PCGs. Two incomplete stop codons, T and A, were exhibited in *nad2* and *nad4*, respectively. As previously reported, incomplete stop codons tend to appear with a high frequency in metazoan mitotic genomes and can be modified by the polyadenylation after transcription (Dreyer and Steiner 2006).

**Table 3.** The genome composition and the AT and GC skews of *H. japonica*.

| Composition | Length(bp) | A% | T% | G% | C% | AT% | GC% | AT-Skew | GC-Skew |
| --- | --- | --- | --- | --- | --- | --- | --- | --- | --- |
| Mitogenome | 15,980 | 36.62 | 36.9 | 8.98 | 17.5 | 73.52 | 26.48 | -0.0079 | 0.322 |
| PCGs | 11,057 | 30.61 | 41.22 | 13.82 | 14.34 | 71.84 | 28.16 | -0.148 | -0.018 |
| Third position | 3,685 | 27.43 | 44.71 | 13.38 | 14.46 | 72.14 | 27.84 | -0.240 | -0.039 |
| tRNAs | 1,437 | 40.57 | 37.58 | 12.53 | 9.32 | 78.15 | 21.85 | 0.038 | 0.147 |
| rRNAs | 1,809 | 39.03 | 39.19 | 14.37 | 7.41 | 78.22 | 21.78 | -0.022 | 0.320 |

The mitogenomes of metazoans are always biased towards A and T, which directly enables the bias of their corresponding amino acids. The AT and GC skews of *H. japonica* PCGs are -0.148 and -0.018, respectively (Table 3), indicating that encoded genes have a significant bias in the use of T and a light bias in the use of C. Furthermore, synonymous codons with A and T at the third position are applied much more frequently than other codons. This phenomenon reveals the A and T preferences at the third position, which resembles those in many metazoans (Chai *et al*. 2012; Zhang *et al*. 2017; Sun *et al*. 2018b). The relative synonymous codon usages (RSCUs) of *H. japonica* PCGs are shown in Fig. 2. 3,687 amino acids were totally encoded in the mitogenome of *H. japonica*. Here, serine (16.59%) and tryptophan (1.64%) owned the highest and lowest use frequency, respectively. Serine rather than leucine was the most used, and AGA (Ser) played an important role in codon usage. As shown, the most commonly used six codons were TTA (Leu), AGA (Ser), TCT (Ser), GCT (Ala), CCT (Pro), and Gly (GGA). Comparative analysis of amino acid composition based on RSCU and PCGs indicated that the codon usage of *H. japonica* was not conservative, which was different from that of most Brachyura species, for example, *Grapsus albolineatus* and *Metopaulias depressus* (Lu *et al*. 2022; Rodriguez-Pilco *et al*. 2022).

**Fig. 2.**
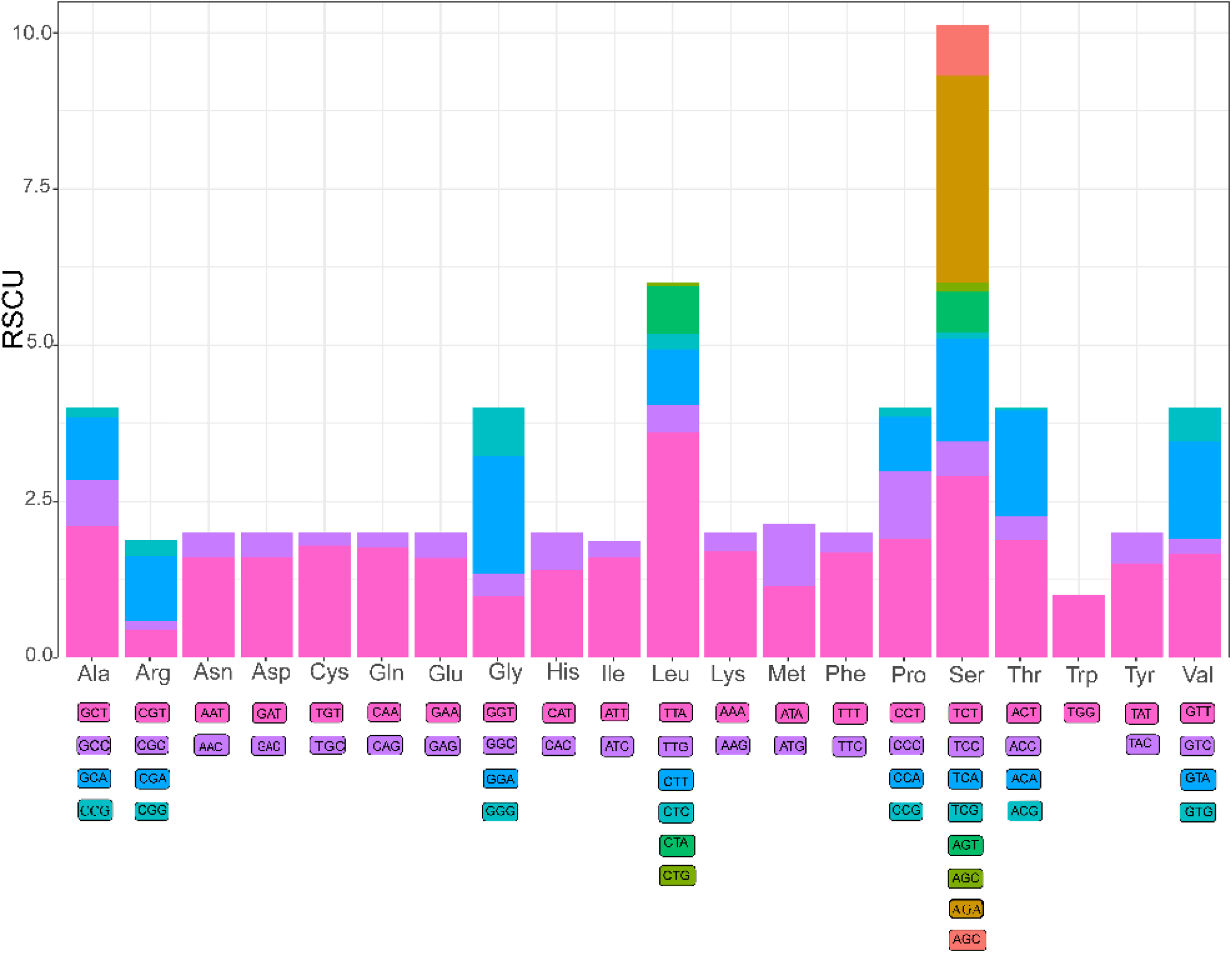
Relative synonymous codon usage (RSCU) of *H. japonica* mitochondrial PCGs. Numbers to the left mean the RSCU values. The X axis represenst codon families.

### 3.3 Transfer RNA and ribosomal RNA genes

Like most crabs, *H. japonica* mitogenome contains 22 tRNA genes and 2 rRNA genes. The size of tRNA genes is 1, 437 bp, 8.99% of the mitogenome (Table 3). The lengths of the 22 tRNA genes range from 64 to 69 bp (Table 2). The AT content and GC content of tRNA regions are 78.15% and 21.85%, respectively. Both of the skews of AT and GC are positive, and their values are 0.038 and 0.147, separately, showing that the use of A is slightly biased, and the use of G is apparently biased.

The two rRNA genes are *rrnL* and *rrnS*, with lengths of 1, 045 and 764 bp, separately (Table 3). Two genes are identified on the L-strand, and account for 11.32% of the total length of *H. japonica* mitogenome. These two genes are separated by gene *trnV(gta)* (Table 2), a feature often occurs in the many Brachyura mitogenomes (Lu *et al*. 2022; Pang *et al*. 2022; Rodriguez-Pilco *et al*. 2022). The AT content of rRNAs is calculated to be 78.22% and the GC content is 21.78%. AT and GC skews are -0.022 and 0.32, respectively, indicating significant biases in the use of T and G.

### 3.4 Phylogenetic relationship

Phylogenetic position of *H. japonica* was investigated with the mitogenome sequences of 40 Brachyuran species and two outgroups (Fig. 3). The results showed that *H. japonica* had the closest relationship with *Pyrhila pisum*, which belongs to the family Leucosiidae. The 40 Brachyuran species and *H. japonica* were separated into two clades. *Calappa Bilineata* formed a seperate clade separately, and the remain 39 species formed the second clade. It is easy to be seen that two families in the superfamily Leucosioidea, Calappidae and Leucosiidae, belonged to two different clades in the phylogenetic tree. Meanwhile *H. japonica* and *Orithyia sinica* in the same superfamily Dorippoidea were not shown to have a close phylogenetic relationship. This proposes challenges for the traditional classification based on morphological data. Although relationships within the superfamily have not been clearly corrected, it can also be seen that the family Dorippidae has a clear trend of close relationship with the family Leucosiidae and then the family Matutidae. This provides important information for the phylogeny of Dorippidae within infraorder Brachyura.

**Fig. 3.**
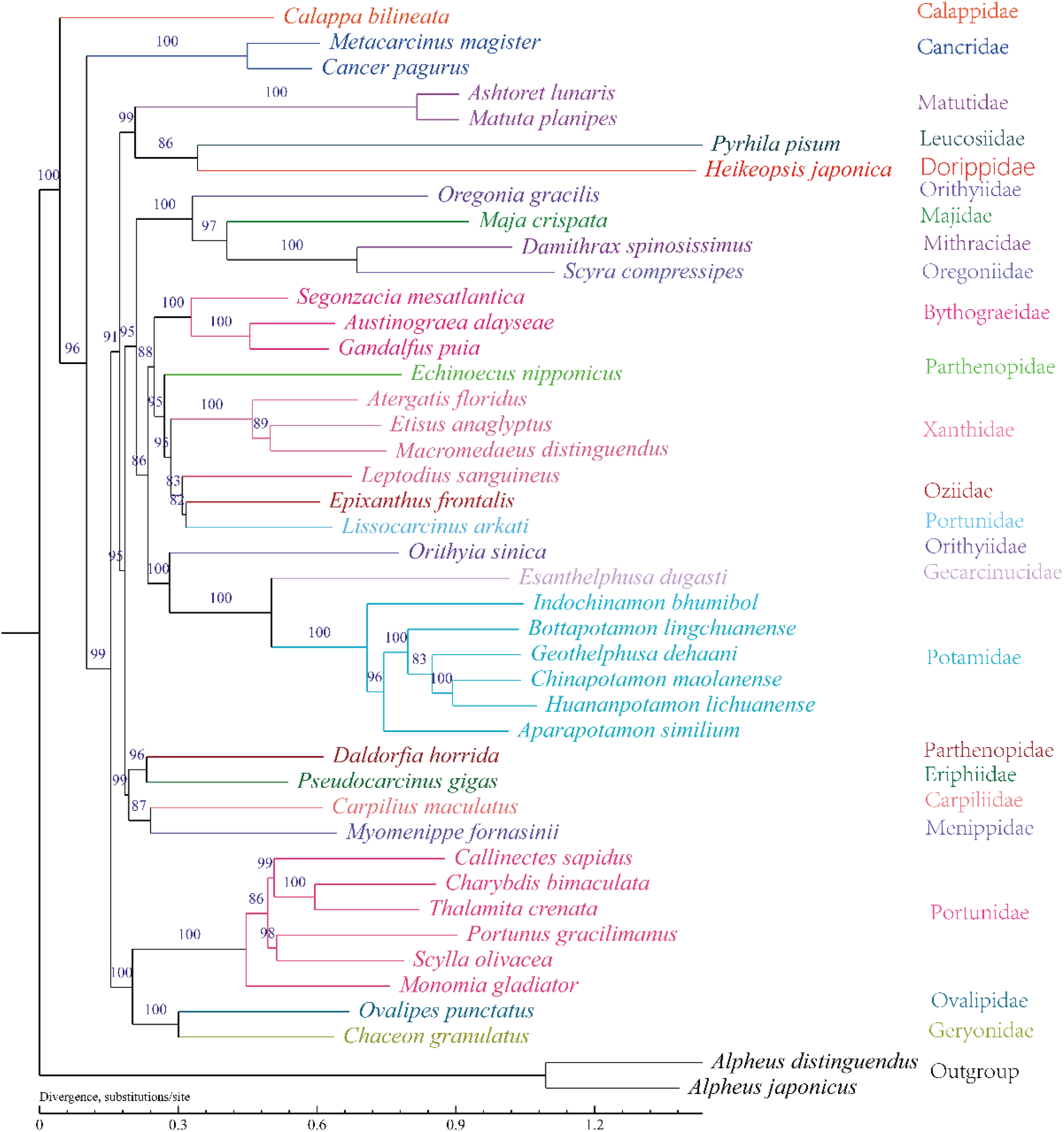
Phylogenetic tree derived from Maximum likelihood (ML) analysis based on 13 mitochondrial PCGs. The right of the tree is the family to which the species belongs. Above the nodes, the number is the bootstrap proportion (BP) value.

### 3.5 Natural selection analysis

To explore the natural selection pressure of *H. japonica* in Brachyuran species, we calculated the ω ratios of 13 mitochondrial PCGs. ω ratio of *H. japonica* calculated under M0 was 0.06422 (Table 4), revealing that the genes were under constrained selection pressure. The ω value of foreground branch under M2 was shown to be 33.41128, much higher than that of the background branch, 0.06373. Meanwhile, the ω ratio of foreground lineage (ω2= 33.41128) under M1 was significantly greater than 1, which represents the strong positive selection of *H. japonica* in Brachyura. The positively selected sites in *H. japonica* were further examined with branch-site models. A total of 31 residues in 12 genes (PCGs expect for *nad6*), were determined as the positively selected amino acid mutation sites, all of which owned high BEB values over 95%. This suggests that the 12 genes are under the heavy force of positive selection (Table 5). The PCGs in mitogenome occupy an important status in the formation of ATP in biological oxidation reaction (Yang *et al*. 2019). These 12 genes containing potential positive selection residues are important components of the aerobic respiration genes and may have essential functions.

**Table 4.** CodeML analysis of the purification selection pressure on the genes of *H. japonica*. M0, M1 and M2 represent one-ratio, free-ratio and two-ratio models, respectively.

| Trees | Models | lnL | Parameter estimates | Model compared | 2ΔL | LRT P-value |
| --- | --- | --- | --- | --- | --- | --- |
| Branch models |  |  |  |  |  |  |
| ML tree | M0 | -229992.3853 | $\omega=0.06422$ | | | |
| | M2 | -229951.0333 | $\omega_0=0.06373$<br>$\omega_1=33.41128$ | M2 vs. M0 | 82.704112 | 0 |
|  | M1 | -229156.5649 |  | M1 vs. M0 | 1,671.64 | 0 |
| Branch-site models |  |  |  |  |  |  |
| ML tree | Null model | -223520.1695 | $p_0=0.00000$;<br>$p_1=0.00000$;<br>$p_{2a}=0.86857$;<br>$p_{2b}=0.13143$<br>$\omega_0=0.04733$;<br>$\omega_1=1.00000$;<br>$\omega_{2a}=1.00000$;<br>$\omega_{2b}=1.00000$ | | | |
| | Model A | -223508.3895 | $p_0=0.76612$;<br>$p_1=0.11577$;<br>$p_{2a}=0.10260$;<br>$p_{2b}=0.01550$<br>$\omega_0=0.04732$;<br>$\omega_1=1.00000$;<br>$\omega_{2a}=999.00000$;<br>$\omega_{2b}=999.00000$ | Model A vs. null model | 23.559876 | 0.000001211 |

**Table 5.** Potentially possible selection sites in *H. japonica*.

| Gene | Codon | Amino acid | BEB values |
| --- | --- | --- | --- |
| *atp6* | 18 | S | 0.960* |
| *atp6* | 31 | I | 0.975* |
| *atp6* | 103 | A | 0.955* |
| *atp6* | 123 | H | 0.959* |
| *atp6* | 175 | G | 0.957* |
| *atp8* | 254 | P | 0.982* |
| *cox1* | 456 | S | 0.953* |
| *cox2* | 776 | T | 0.957* |
| *cox2* | 819 | A | 0.963* |
| *cox2* | 962 | L | 0.985* |
| *cox2* | 989 | S | 0.956* |
| *cox2* | 1000 | G | 0.998** |
| *cox3* | 1159 | G | 0.956* |
| *cox3* | 1184 | A | 0.959* |
| *cox3* | 1208 | S | 0.952* |
| *cob* | 1315 | S | 0.983* |
| *cob* | 1443 | M | 0.991** |
| *nad1* | 1796 | L | 0.992** |
| *nad2* | 1955 | M | 0.953* |
| *nad2* | 2015 | T | 0.970* |
| *nad2* | 2201 | T | 0.997** |
| *nad3* | 2255 | S | 0.952* |
| *nad4* | 2501 | C | 0.962* |
| *nad4* | 2561 | S | 0.954* |
| *nad4* | 2609 | S | 0.967* |
| *nad4* | 2691 | S | 0.963* |
| *nad4* | 2742 | C | 0.987* |
| *nad4L* | 2813 | Y | 1.000** |
| *nad5* | 3068 | S | 0.973* |
| *nad5* | 3100 | S | 0.952* |
| *nad5* | 3131 | H | 0.971* |

\* 0.95 < BEB < 0.99

Complex of NADH dehydrogenase is confirmed to be the largest enzyme complex in biological aerobic respirations, which play the role of active transport of hydrogen ions. Site changes may affect the hydrogen ion transport efficiency, thus affecting the cell metabolic properties (da Fonseca *et al*. 2008). The results showed that a total of 14 potential sites were detected in *nad1-5* and *nad4L* genes (subunits 1-5 and 4L of NADH deshydrogenase). Cytochrome *c* oxidase is responsible for catalyzing the reduction of oxygen terminal, and three PCGs (*cox1-3*) encoded the catalytic core (Luo *et al*. 2008; Yang *et al*. 2019). Nine positively selected sites were totally found in *cox1-3* genes here. The large distribution of the potential selected sites on the *nad* genes as well as *cob* genes provides novel insights into the strong positive selection pressure on *H. japonica* in the infraorder Brachyura.

## 4. Conclusions

In this study, the first species mitogenome of family Dorippidae, *H. japonica*, was sequenced and characterized. It is a closed circular comprising 13 PCGs, 22 tRNAs, and 2 rRNAs. The phylogenetic tree of Brachyuran species showed that *Pyrhila pisum* of the family Leucosiidae is most closely related to *H. japonica*. A total of 31 residues located in 12 PCGs were further detected as positively selected sites. This provides direct evidence that *H. japonica* is under strong positive selection pressure in Brachyura. The results offer meaningful mitogenome information for broadening the knowledge of marine crabs, and propose new insights of the phylogenetic status and positive selection pressure of family Dorippidae within infraorder Brachyura.

## Data availability

The mitogenome reads are available from NCBI with number OQ434093.

## Acknowledgements

The authors thank Tianjin Fisheries Research Institute for sampling and Shanghai BIOZERON Co., Ltd. for Illumina sequencing.

